# Improved LC-MS chromatographic alignment increases the accuracy of label-free quantitative proteomics: Comparison of spectral counting versus ion intensity-based proteomic quantification strategies

**DOI:** 10.1101/111476

**Authors:** Daniel H.J. Ng, Jonathan D. Humphries, Julian N. Selley, Stacey Warwood, David Knight, Adam Byron, Martin J. Humphries

**Affiliations:** Wellcome Trust Centre for Cell-Matrix Research, Faculty of Biology, Medicine and Health, University of Manchester, Manchester M13 9PT, UK; Biological Mass Spectrometry Core Facility, Faculty of Biology, Medicine and Health, University of Manchester, Manchester M13 9PT, UK; Edinburgh Cancer Research UK Centre, Institute of Genetics and Molecular Medicine, University of Edinburgh, Edinburgh EH4 2XR, UK

## Abstract

The ability to provide an unbiased qualitative and quantitative description of the global changes to proteins in a cell or an organism would permit the systems-wide study of complex biological systems. Label-free quantitative shotgun proteomic strategies (including LC-MS ion intensity quantification and spectral counting) are attractive because of their relatively low cost, ease of implementation, and the lack of multiplexing restrictions when comparing multiple samples. Owing to improvements in the resolution and sensitivity of mass spectrometers, and the availability of analytical software packages, protein quantification by LC-MS ion intensity has increased in popularity. Here, we have addressed the importance of chromatographic alignment on protein quantification, and then assessed how spectral counting compares to ion intensity-based proteomic quantification. Using a spiked-in protein strategy, we analysed two situations that commonly arise in the application of proteomics to cell biology: (i) samples with a small number of proteins of differential abundance in a larger non-changing background, and (ii) samples with a larger number of proteins of differential abundance. To perform these assessments on biologically relevant samples, we used isolated integrin adhesion complexes (IACs). Technical replicate analysis of isolated IACs resulted in a range of alignment scores using the Progenesis QI software package and demonstrated that higher LC-MS chromatographic alignment scores increased the precision of protein quantification. Furthermore, implementation of a simple sample batch-running strategy enabled good chromatographic alignment for hundreds of samples over multiple batches. Finally, we applied the sample batch-running strategy and compared quantification by LC-MS ion intensity to spectral counting and found that quantification by LC-MS ion intensity was more accurate and precise. In summary, these results demonstrate that chromatographic alignment is important for precise and accurate protein quantification based on LC-MS ion intensity and accordingly we present a simple sample re-ordering strategy to facilitate improved alignment. These findings are not only relevant to label-free quantification using Progenesis QI but may be useful to the wide range of MS-based quantification strategies that rely on chromatographic alignment.

## Introduction

The aim of functional proteomics is to characterise biological systems in a global and unbiased manner, such that the biological function of proteins can be inferred from their interacting partners, abundance, post-translational modifications and cell localisation [1]. The qualitative and quantitative description of such biological systems allows for systems-wide characterisation of biochemical pathways and biomarker discovery. Historically, proteomic studies involved protein profiling through the use of two-dimensional gel electrophoresis [2]. Current “bottom-up” shotgun mass spectrometry MS-based approaches permit both the identification and quantification of thousands of proteins accurately across different conditions [2,3].

A typical shotgun MS proteomic experiment involves enzymatic digestion of proteins into peptides, followed by high-performance liquid chromatographic (LC) separation of peptides and on-line analysis of the eluting peptides by MS [4]. Peptide samples are commonly vaporised and ionised by electrospray ionisation (ESI). An initial survey scan (MS1) detects the mass-to-charge *(m/z)* ratio and the intensity of precursor peptide ions. Thereafter, signal intensity heuristics are used during data-dependent acquisition (DDA) to select the most abundant precursor peptide ions for collision-induced dissociation. The resulting fragment-ion mass spectrum is measured in a second MS scan (MS2), the overall process being termed tandem mass spectrometry (MS/MS). Finally, bioinformatic tools utilise information from the MS1 and MS2 spectra to identify peptides sequences, from which protein identities are inferred.

Quantification methods for proteins during shotgun MS proteomic experiments fall broadly into two categories: those that incorporate a stable isotope label or those that do not (label-free) [3]. Compared to stable isotope labelling strategies, label-free quantification strategies are attractive because of their relatively straightforward implementation, lack of expensive isotope labels, and the ability to compare any number of conditions [5-9]. There are two commonly used label-free methods: spectral counting and quantification by LC-MS ion intensity [3,5,7,10]. Spectral counting is performed at the MS/MS (fragment ion) level [11,12] with the assumption that all precursor peptides are selected by DDA and are detected by MS/MS: in reality, only a subset of the precursor peptides ions are selected, which biases quantification towards the most abundant peptides. In contrast, LC-MS ion intensity quantification typically involves the integration of the area under the chromatographic peak, or volume of the peak, in the time, m/z and intensity dimensions as a direct measure of peptide abundance [13-15]. Direct comparison of corresponding peptide chromatographic peak volumes across different samples permits the relative quantification of peptides. However, this is complicated by chromatographic drifts that arise during the LC separation process [16,17]. To this end, several groups, both academic and commercial, have developed warping algorithms to reduce shifts in the retention time axis such that corresponding peptide features across multiple conditions can be aligned [9,16,17-20]. This process is important because it facilitates consistent peak picking across multiple conditions, enables appropriate normalisation of data, reduces complications in assigning peptide identifications from MS/MS spectra, and allows the direct comparison of peptide features across multiple conditions [8,10]. Although several studies have examined algorithms used for chromatographic alignment, few studies have addressed the impact of chromatographic alignment on the quantification of biologically relevant protein samples. Furthermore, in light of recent improvements to mass spectrometers and analysis software, it is important to assess and compare the label-free relative quantification strategies, LC-MS ion intensity quantification and spectral counting.

In this study, we have addressed the importance of chromatographic alignment on LC-MS ion intensity-based quantification. By comparing technical replicates with a range of chromatographic alignment scores, we find that chromatographic alignment correlates with the precision of protein quantification. In addition, we developed a simple sample batch-running (blocking) strategy to improve chromatographic alignment by minimising chromatographic drifts and elution profile changes. In combination with peptide spiking, blocking enabled good chromatographic alignment, which permitted protein quantification by LC-MS ion intensity that was more precise and accurate than spectral counting. These analyses were performed in experimental set-ups mimicking two biologically relevant conditions: a few changing proteins in a complex protein environment, and a large number of proteins changing simultaneously. Therefore, this study highlights the caveats to protein quantification by LC-MS ion intensity and will help to direct biologists in their choice of label-free quantification strategy.

## Experimental procedures

### Materials

The protein standard (Dionex, Thermo Fisher Scientific) contained a tryptic digest of carboxymethylated peptides from six non-human proteins [cytochrome c (*Bos taurus*), lysozyme (*Gallus gallus domesticus*), alcohol dehydrogenase (*Saccharomyces cerevisiae*), bovine serum albumin (B. *taurus*), apo-transferrin (B. *taurus*), β-galactosidase (*Escherichia coli*)].

### Preparation of a complex mixture of proteins

The moderately complex mixture of proteins consisted of isolated cell adhesion complexes [21-23] and was selected to represent a biologically relevant level of complexity from a well-studied biological fraction of a cell. Briefly, human foreskin fibroblasts (HFFs) were permitted to interact with the extracellular matrix ligand fibronectin to allow integrin adhesion complexes (IACs) to form on the basal surface of the cells. Cells were cross-linked with 6 mM DTBP for 3 min and quenched with Tris-HCl, pH 8.5, followed by sonication for 2.5 min to lyse cells to remove the bulk of the cell. The integrin adhesion complexes (IACs) that remained adherent to the dish were collected in reducing Laemmli sample buffer [250 mM Tris-HCl, 40% (w/v) glycerol, 8% (w/v) sodium dodecyl sulfate (SDS), 0.02% (w/v) bromophenol blue and 10% (v/v) β-mercaptoethanol] and fractionated by SDS-PAGE. In-gel proteolytic digestion was carried out as described by Horton et al. [22]. Briefly, gel lanes were cut into five slices, and each slice cut into ~1 mm^3^ pieces. Gel pieces were destained with 50% (v/v) acetonitrile in 12.5 mM NH4HCO3, dehydrated with acetonitrile, reduced in 10 mM dithiothreitol, alkylated in 55 mM iodoacetamide, dehydrated, and proteins were digested with trypsin (12.5 ng/μl). Peptides were collected in one wash of 99.8% (v/v) acetonitrile and 0.2% (v/v) formic acid and one wash of 50% (v/v) acetonitrile and 0.1% (v/v) formic acid. Peptides were desalted using R3 beads (Applied Biosystems).

### Dilution of protein standard in a constant complex IAC protein background

The protein standard dilution series was carried out as indicated in Fig. 3A, resulting in five moderately complex IAC protein fractions spiked with 4-fold decreasing concentrations of protein standard (40 fmol, 10 fmol, 2.5 fmol, 0.625 fmol and 0.157 fmol), and an additional IAC fraction without spiking was used as a blank. Triplicates of each sample were analysed by LC-MS/MS. Data normalisation was carried out assuming the majority of peptides across conditions were not changing. For LC-MS ion intensity quantification, a global normalisation factor was used; for spectral counting, the unweighted spectral counts were divided by the total number of spectra observed in the entire sample.

### Dilution of the IAC mixture of complex proteins in a constant protein standard

The dilution experiment was carried out as indicated in Fig. 3B, resulting in six dilutions of the moderately complex IAC protein background (no dilution, 1/2, 1/4, 1/8, 1/16 and 1/32). Fifty fmol of protein standard was spiked into each IAC fraction and triplicates of each samples analysed by LC-MS/MS. Data normalisation was carried out assuming that the six-protein mix standard peptides were not changing.

### MS data acquisition and analysis

LC-MS/MS was carried out using an UltiMate 3000 Rapid Separation LC (Dionex) coupled to an Orbitrap Elite (Thermo Fisher Scientific) mass spectrometer. Peptides were separated in an analytical column (250 mm × 75 μm i.d, 1.7 μμm BEH C18; Waters) using an increasing acetonitrile gradient, with a starting mixture of 8% solution A (0.1% formic acid in acetonitrile) and 92% solution B (0.1% formic acid in water) to 33% solution B in 44 min at 300 nL min^−1^. Peptides were selected for fragmentation automatically by data-dependent analysis (DDA).

Peak list files were searched against a modified version of UniProt/Swiss-Prot database (biomols_custom-uniprot_sprot database, version 03_3_24, release date 05-2011, containing 528, 278 sequences) using an in-house Mascot server (version 2.2.06, Matrix Science). Variable modifications were set for oxidation of methionine, carbamidomethylation of cysteine and carboxymethylation of cysteine. Maximum missed cleavages for tryptic peptides was set to one. Only monoisotopic precursor ions that were doubly or triply charged were considered.

### MS data processing

For LC-MS ion intensity quantification, profile RAW files were imported into Progenesis QI software (Nonlinear Dynamics, version 4.1.4832.42146; http://www.nonlinear.com/progenesis/qi/). Alignment of chromatograms was carried out using the automatic alignment algorithm, followed by manual validation and adjustment of the aligned chromatograms. All features were used for peptide identifications.

For spectral counting, Mascot Daemon was used to process batches of RAW files by creating peak lists using a search script (extract_msn; Thermo). Peak lists were searched in Mascot. Results were loaded in Scaffold (Proteome Software, version 3.6.5), and peptide and protein identification thresholds were set to 95% and 99% confidence, respectively. Bovine serum albumin (BSA) was present in the complex protein sample and protein standard, and was excluded from spectral counting analysis.

Limit of detection (LOD) and limit of blank (LOB) were calculated as follows [24]:

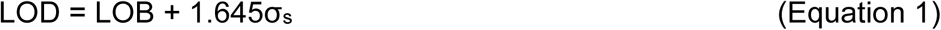

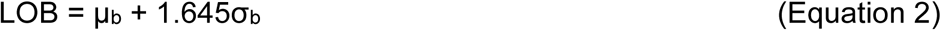

where σ_s_ = standard deviation of low concentration sample, μ_b_ = mean of blank, and σ_b_ = standard deviation of blank.

### Hierarchical clustering

The log_10_(average protein abundance) was calculated for LC-MS ion intensity quantification and spectral counts. To ensure that starting points for data were the same while gradients were unchanged, data was translated in the y-axis such that all data starting points were 1. Data was loaded in MultiExperiment Viewer (TM4 Microarray Software Suite) and hierarchically clustered using Euclidean distance as the distance metric and average linkage as the linkage criteria.

## Results

### Effect of LC-MS chromatographic alignment on ion intensity-based protein quantification

To investigate the effect of chromatographic alignment on protein quantification, we analysed data consisting of several repeat LC-MS/MS analyses of the same complex mixture of integrin adhesion complex (IAC) proteins [21-23]. Technical replicates consisting of identical peptide runs were analysed using Progenesis QI. The Progenesis QI software platform (http://www.nonlinear.com/progenesis/qi/) functions by the selection of a reference LC-MS sample dataset and the alignment of other LC-MS sample runs to this reference. In this way, Progenesis QI can be used to compensate for between-run variation in the LC separation retention times, and the objectively determined alignment score provides a measure of the similarity and quality of the LC-MS runs within an experiment.

For this analysis, proteins in replicate LC-MS runs of IACs with a range of Progenesis QI LC-MS chromatographic alignment scores were identified and quantified (Fig. 1). Although the chromatographic alignment scores are dependent on the degree of overlap between features, and misalignment of conflicting features may still yield positive alignment scores, we used these scores as a qualitative measure of the LC-MS alignment, along with visual interpretation to determine alignment success. Theoretically, quantitative analysis through the Progenesis QI LC-MS pipeline should result in proteins with an expected fold change in abundance of one between technical replicates. Indeed, the majority of proteins (at least 65%) across all the datasets were within a 1.5-fold change. As the chromatographic alignment scores of samples decreased, there was a larger spread of fold changes (Fig. 1). Moreover, the relative number of proteins detected in only one sample increased as shown by the increase in the percentage of infinite points (Fig. 1). We observed a notable increase in the spread of the distribution of fold changes between runs at approximately less than 90% alignment. From these analyses, we conclude that chromatographic alignment scores should be at least >80%, and ideally >90%, to minimise variation and improve precision in quantification. Interestingly, it was observed that poor chromatographic alignment arose when there were several features with a different order of elution (elution profile) from the reference run. In contrast, features having the same elution profile as the reference run that displayed chromatographic drifts aligned well after applying the pre-set warping algorithm, indicating that while the warping algorithm deals well with chromatographic drifts, it is unable to deal with changes in the chromatographic elution profile.

**Fig 1.**
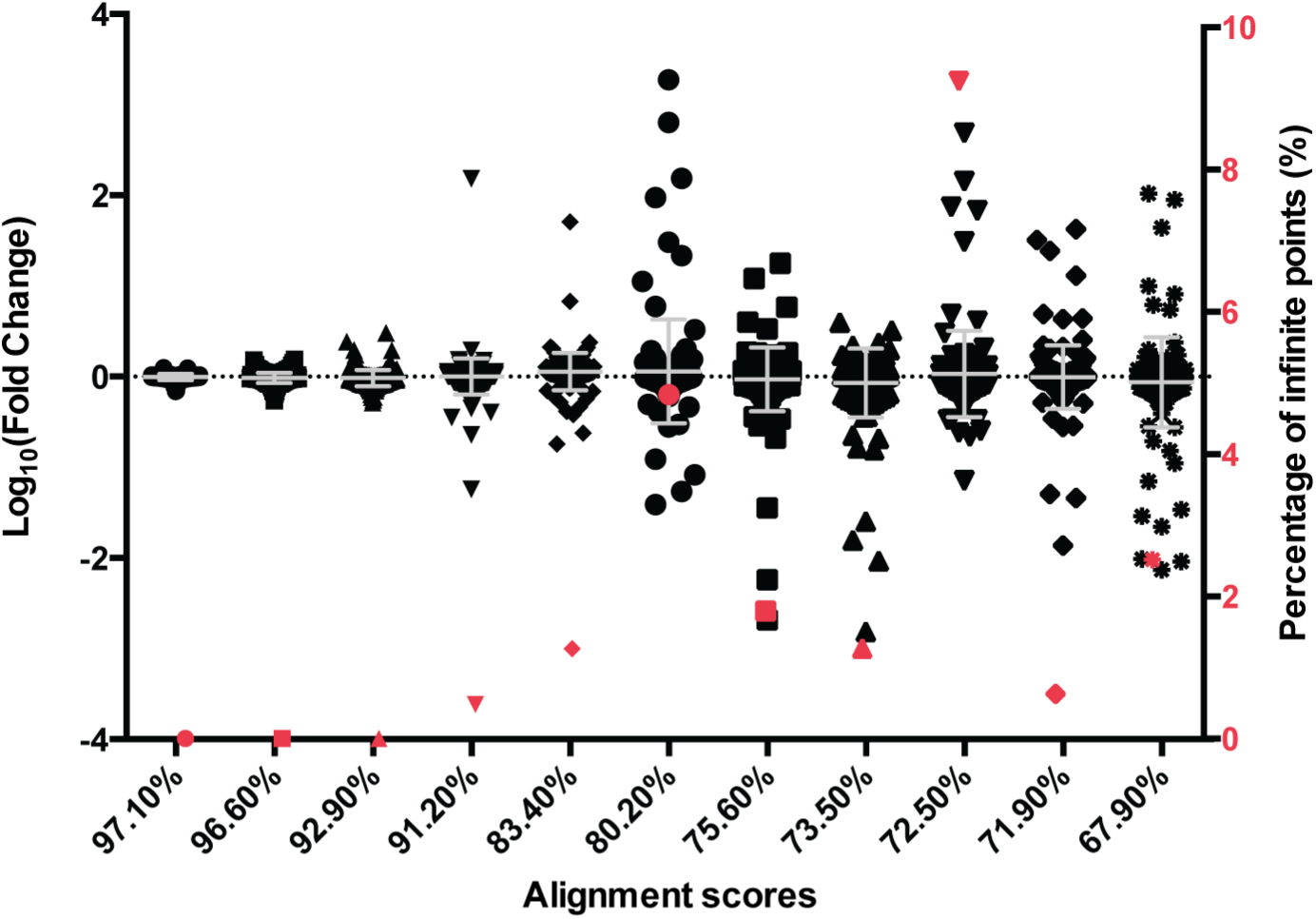
Effect of LC-MS chromatographic alignment on protein quantification. Technical replicates were compared and protein fold changes (left y-axis, black points) and number of infinite points (right y-axis, red points) were plotted against the Progenesis QI LC-MS alignment scores.

### Strategy to improve LC-MS chromatographic alignment

Chromatographic drifts and changes to the sequence of the chromatographic elution profile may occur because of subtle changes in chemical environment or performance of the LC instrument over time owing to the interaction of the LC column with samples. In general, we noticed that technical replicates with poor alignment scores were usually run far apart in time, and usually in different sample batches, interspersed with washes and calibration runs. To minimise chromatographic drifts and to ensure that peptides maintain the sequence of their elution profile over the samples being compared, we hypothesised that samples to be directly compared should be analysed in the same sample batch (block), close together in time (Fig. 2). Additionally, a pooled reference sample, comprising a small proportion of peptides from all samples to be analysed, was used as a reference for alignment or to equilibrate the LC column. In experiments where fractionation is carried out, samples across one fraction are analysed in the same block (Fig. 2). To test the performance of this strategy, Progenesis QI chromatographic alignment scores were assessed for a large-scale experiment (208 samples over 16 batches). These analyses revealed that good alignment (>85%) was achieved by this sample blocking strategy for all samples.

**Fig 2.**
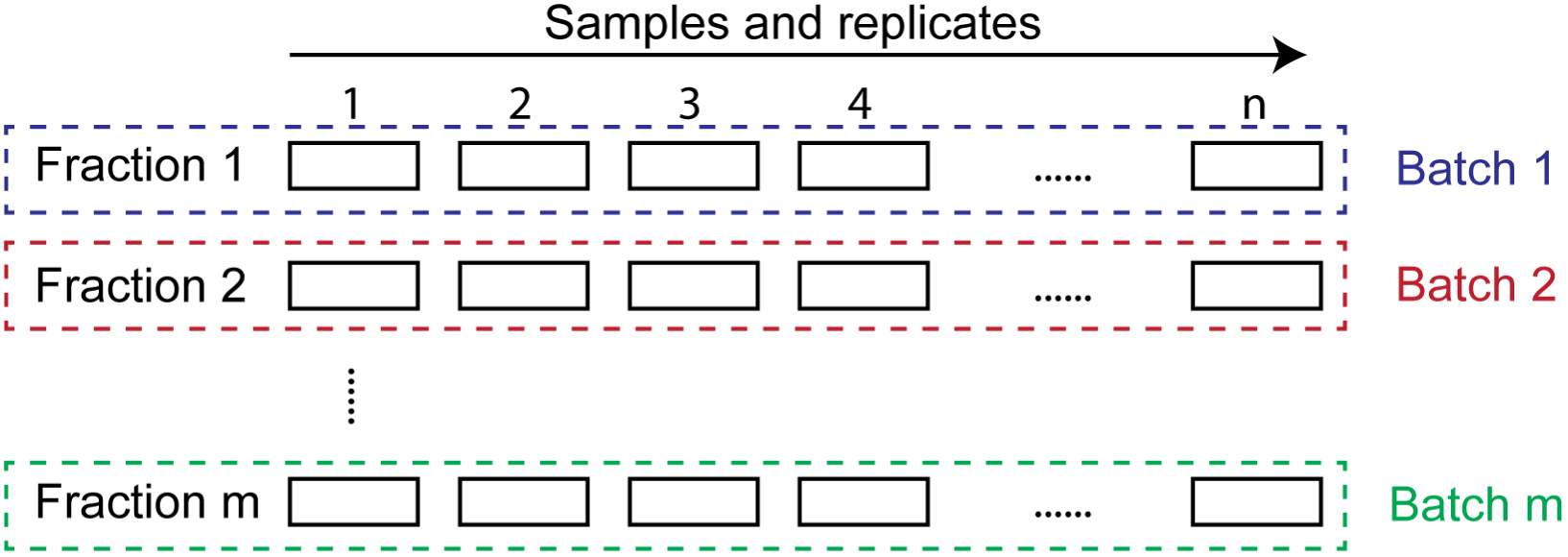
Strategy to improve chromatographic alignment. Samples and replicates to be compared are analysed in the same sample batch. Separate fractions are analysed in different sample batches.

### Experimental set-up to test label-free quantification methods

Having established that chromatographic alignment affects the precision of LC ion intensity quantification and that alignment can be improved by sample blocking, we wanted to test LC-MS ion intensity quantification on biologically relevant samples and benchmark it against another commonly used label-free quantification strategy, spectral counting. We devised an experimental set-up to mimic two types of biologically relevant conditions: a few changing proteins in a moderately complex IAC protein environment, and a large number of IAC proteins changing simultaneously. To quantify a few changing proteins in a complex protein environment, a moderately complex human IAC protein background was spiked with non-human protein standard at 4-fold decreasing dilutions in triplicate (Fig. 3A). To further determine the ability of label-free quantitative methods to measure the abundance changes of a large number of proteins simultaneously, a moderately complex human IAC protein sample was diluted 2-fold sequentially up to a maximum of 32-fold. The large changes to the majority of peptides prevent the proper alignment of chromatograms, and skew the global normalisation factor; therefore, the protein standard was spiked in at a constant concentration of 50 fmol to provide a reference for chromatographic alignment, and to normalise data.

**Fig 3.**
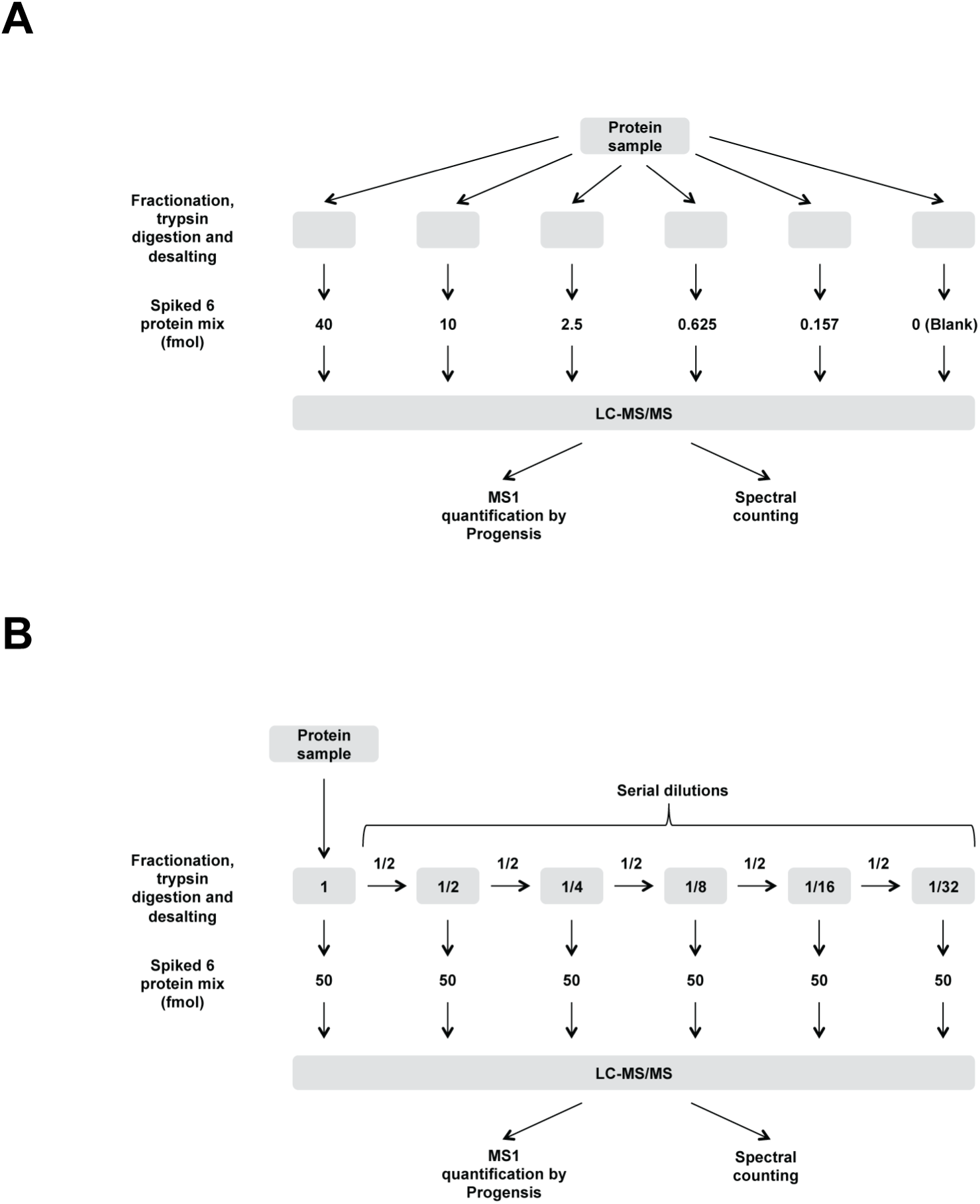
Experimental workflow for the comparison of label-free quantitative MS methods. (A) Complex protein sample of isolated integrin adhesion complex proteins [21-23] was prepared as five fractions and spiked with protein standards at 4-fold reducing concentrations, with an additional sample used as the blank. (B) Complex protein sample of isolated integrin adhesion complex proteins was diluted 2-fold sequentially over a 32-fold range and spiked with 50 fmol of protein standard.

### Quantification of diluted protein standards by LC-MS ion intensity and spectral counting

We first sought to assess the accuracy and precision of protein quantification by LC-MS ion intensity measurements. All samples were determined to have good Progenesis QI alignment scores (>85%). Protein standards showed linear responses between measured protein abundance and protein concentration, above the overall limits of detection (LOD), from 2.5 to 40 fmol (Fig. 4A and Table 1; R^2^ ≥ 0.987) with similar gradients (Fig. 4A and Table 1; 1.17 ≤ gradient ≤ 1.26), indicating that protein quantification changed at almost the same rate upon dilution. The calculated fold change for each protein (Table 1; 4.94 ≤ fold change ≤ 5.74) slightly overestimated the fold change but was in general agreement with the expected 4-fold change. To assess the precision of protein quantification by LC-MS ion intensity, coefficients of variation (CV) for protein quantification were calculated and showed that while CVs increased with decreasing concentration of protein standard, CVs were less than 20% in all the conditions tested (Fig 4B). Taken together, these results indicate that changes to protein abundance in a moderately complex IAC background can be quantified precisely and relatively accurately by LC-MS ion intensities.

**Table 1.** Protein standard dilution series variables for protein quantification by LC-MS ion intensity.

| **Proteins** | **Gradient** | **$R^2$** | **Fold change** | **LOD (fmol)** |
| --- | --- | --- | --- | --- |
| Cytochrome c | 1.17 | 0.997 | 5.03 | 0.72 |
| Lysozyme C | 1.15 | 0.987 | 4.94 | 1.07 |
| Alcohol dehydrogenase 1 | 1.19 | 0.987 | 5.19 | 3.48 |
| Serotransferrin | 1.26 | 0.991 | 5.74 | 1.16 |
| Beta-galactosidase | 1.24 | 0.990 | 5.59 | 1.28 |
| Serum albumin (Carboxymethyl) | 1.20 | 0.994 | 5.25 | 0.52 |

\* Variables for standard curve were calculated using LC-MS ion intensity readings from 2.5 to 40 fmol.

**Fig 4.**
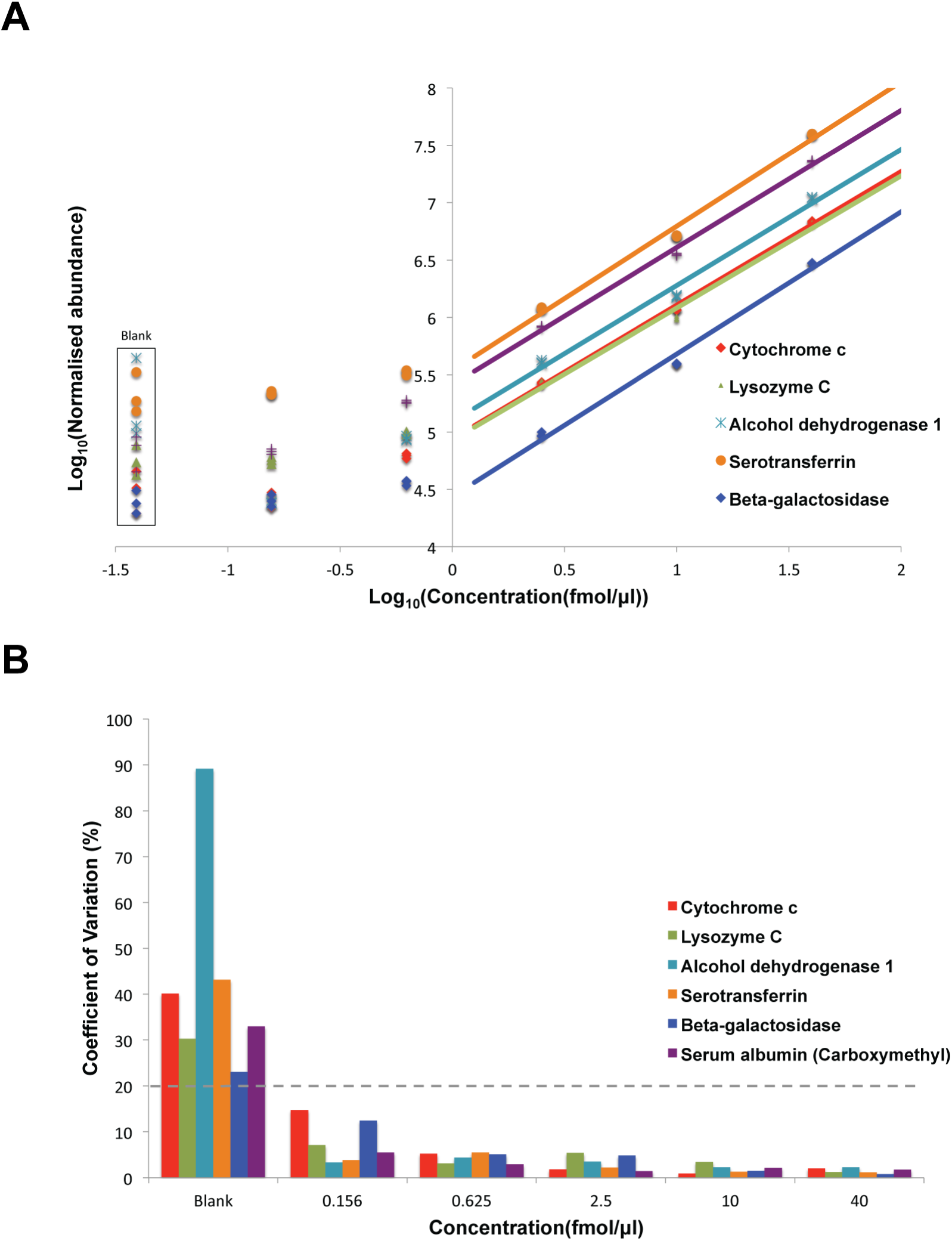
Quantification of protein standard dilution series by LC-MS ion intensity. (A) Standard curve of log_10_(normalised abundance) as measured by LC-MS ion intensities against log_10_(concentration) of protein standard. (B) Frequency distribution plot of LC-MS ion intensity CVs of protein standards. Dotted line indicates 20% CV threshold.

Next, we sought to assess the accuracy and precision of protein quantification by spectral counting. Spectral counts of proteins responded linearly with protein concentration (Fig. 5A and Table 2; 0.648 ≤ R^2^ ≤ 0.969); however, gradients were different for each protein (Fig. 5A and Table 2; 0.15 ≤ gradient ≤ 0.58), indicating that protein quantification changed at different rates for each protein upon dilution. For all proteins, spectral counting underestimated the expected 4-fold change in protein abundance (Table 2; 1.23 ≤ fold change ≤ 2.22). To assess the precision of protein quantification by spectral counting, protein CVs were calculated, and similarly to protein quantification by LC-MS ion intensity, CVs increased with decreasing concentration of protein standard. However, while all proteins at 40 fmol were below a 20% CV threshold, only two out of five proteins at 2.5 fmol had CVs less than the CV threshold (Fig. 5B). Moreover, at each concentration, protein CVs for spectral counting were generally higher than protein quantification by LC-MS ion intensity (Figs 4B and 5B). In summary, spectral counting correlated linearly to changes of protein abundance in a moderately complex IAC background; however, this method underestimated the expected fold change and displayed higher variability, particularly at low protein concentrations compared to quantification using LC-MS ion intensity.

**Table 2.** Protein standard dilution series for protein quantification by spectral counting.

| **Proteins** | **Gradient** | **$R^2$** | **Fold change** |
| --- | --- | --- | --- |
| Cytochrome c | 0.29 | 0.845 | 1.49 |
| Lysozyme C | 0.37 | 0.866 | 1.71 |
| Alcohol dehydrogenase 1 | 0.15 | 0.648 | 1.23 |
| Serotransferrin | 0.42 | 0.969 | 1.80 |
| Beta-galactosidase | 0.58 | 0.903 | 2.22 |

\* Variables for standard curve were calculated using spectral count readings from 2.5 to 40 fmol.

**Fig 5.**
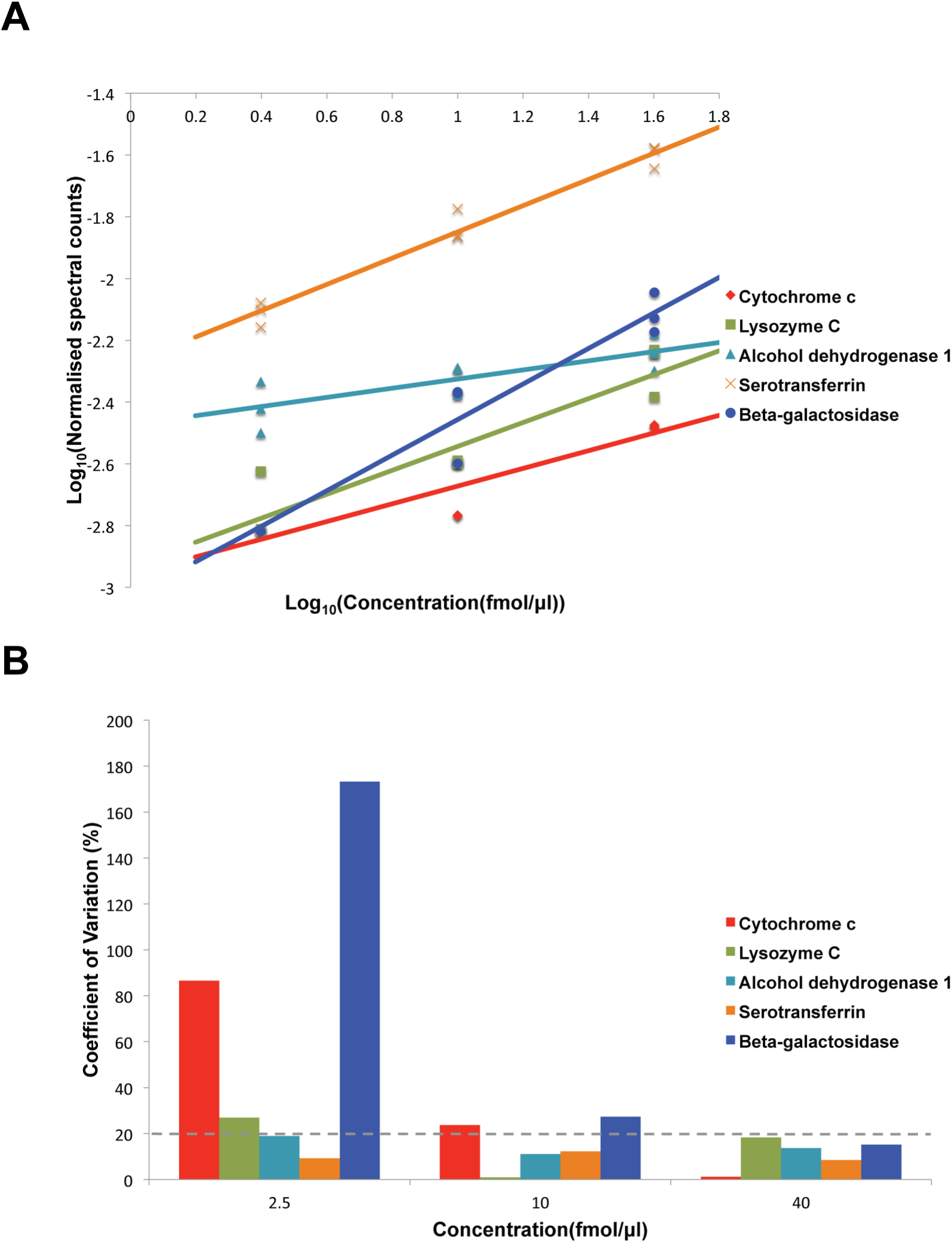
Quantification of protein standard dilution series by spectral counting. (A) Standard curve of log_10_(spectral counts) against log_10_(concentration) of protein standard. (B) Frequency distribution plot of spectral count CVs of protein standard. Dotted line indicates 20% CV threshold. Fig 6.

To compare LC-MS ion intensity quantification to spectral counting directly, the scaled log_10_ (normalised abundance) from ion intensity measurements were plotted against the scaled log_10_ (spectral counts) for each protein (Fig. 6). Protein standards responded linearly but deviated from the ideal regression (grey line), which assumed that both LC-MS ion intensity quantification and spectral counting performed equally well. At 10 fmol, quantification of protein standard was measured to be around the expected value for LC-MS ion intensity. At 40 fmol, quantification of protein standard was measured to be more than the expected value. In contrast, all values for spectral counting at 10 and 40 fmol were less than the expected value. Strikingly, large variances were observed along the x-axis and not the y-axis, which is in agreement with the finding that CVs were larger for spectral counting than with LC-MS ion intensity quantification. Taken together, these results indicate that whilst spectral counting does correlate with protein abundance, LC-MS ion intensity quantification is more accurate and precise in determining relative protein changes particularly at lower concentrations near to that of the LOD.

**Fig 6.**
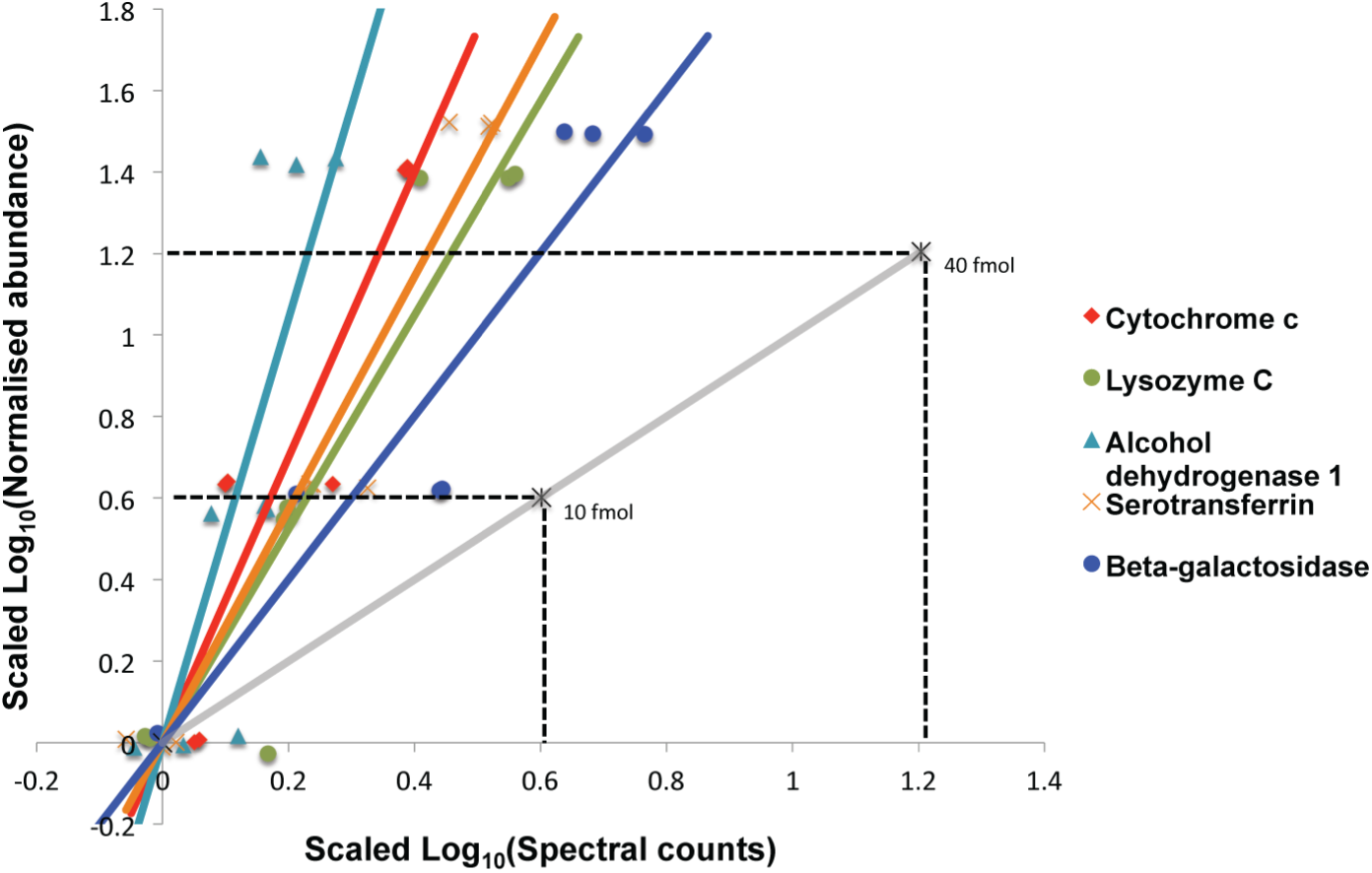
Comparison of protein quantification by LC-MS ion intensity and spectral counting. Direct comparison of LC-MS ion intensity quantification and spectral counting by plotting scaled log_10_(normalised abundance) against scaled log_10_(spectral counts). Dotted lines indicate the expected value at that particular concentration. Grey line indicates the theoretical comparison.

### Quantification of a diluted complex protein mix by LC-MS ion intensity and spectral counting

Unsupervised hierarchical clustering was used to provide an unbiased overview of the pattern of abundance changes to proteins (Figs 3B and 7). Protein quantification using LC-MS ion intensity data revealed two distinct clusters: 1) proteins that did not change upon dilution, and 2) proteins that decreased upon dilution (Fig. 7A). Proteins in the non-changing spiked-in protein standard were found in cluster 1, and proteins in the diluted complex protein background were found in cluster 2. The calculated median fold change of proteins from the protein standard and in cluster 2 were 0.91 and 1.98-fold, respectively, which were near that of the expected fold change of 1 and 2-fold, respectively (Fig. 7B). To determine the precision of protein quantification by LC-MS ion intensity, CVs were calculated for proteins in cluster 2 (Fig. 7C). Once again, CVs were larger for increased dilutions of proteins. At zero dilution, 1/2 and 1/4 dilutions, the majority of proteins fell within a 20% CV threshold (~95%, 95% and 86%, respectively), whereas at 1/8, 1/16 and 1/32 dilutions, fewer proteins fell within the 20% CV threshold (~43%, 34% and 52%, respectively) (Fig. 7C). These results indicate that LC-MS ion intensity-based measurements can accurately and precisely quantify abundance changes for a large number of proteins in a moderately complex protein sample.

**Fig 7.**
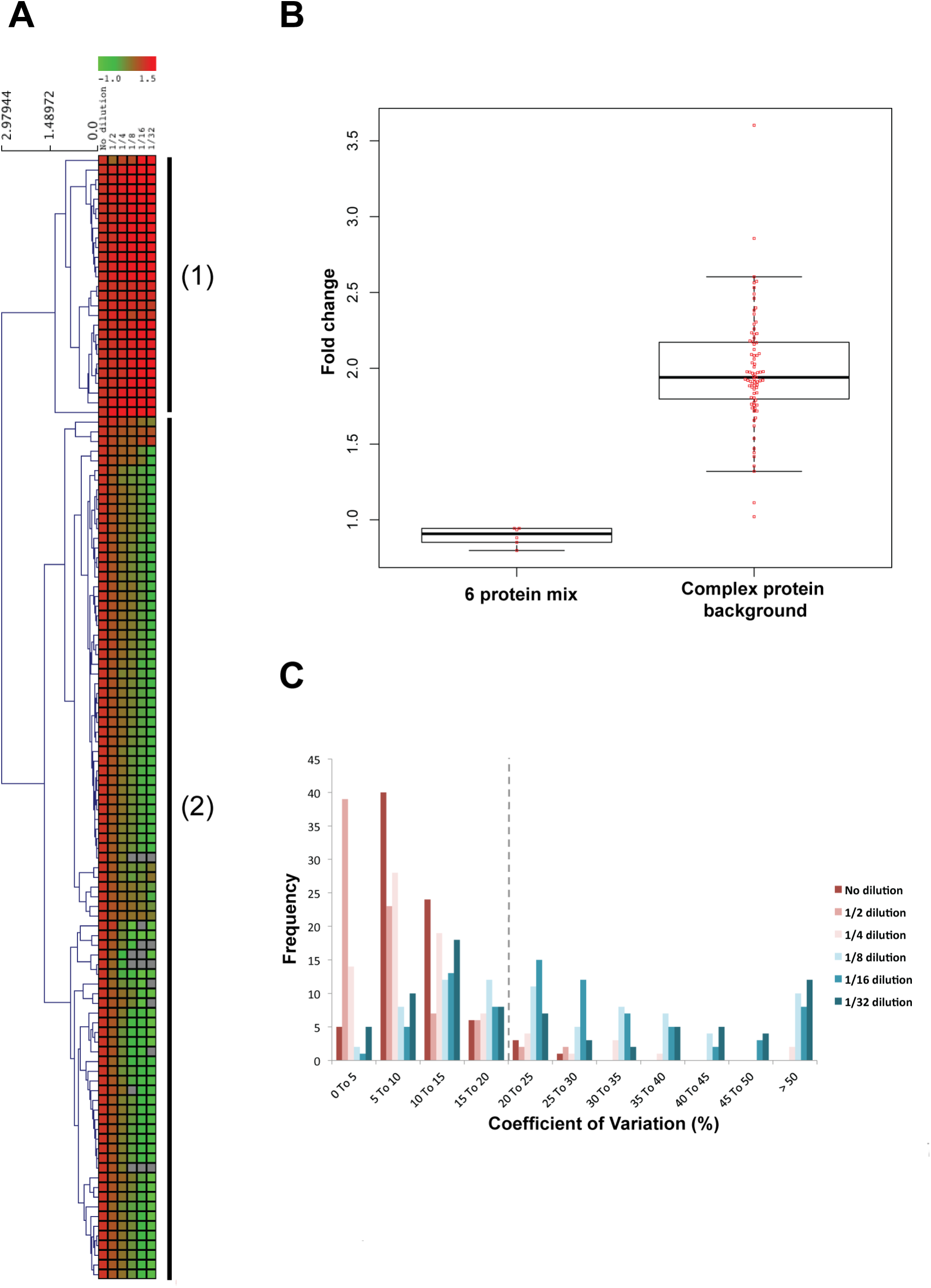
Quantitative analysis of changes to a complex protein mixture dilution series by LC-MS ion intensity. (A) Heat map and dendrogram displaying the Euclidean distance-based hierarchical clustering of the scaled log_10_(average normalised abundance) over six dilutions (no dilution to 1/32 dilution). Two main clusters were observed: 1) protein with abundances that did not change, and 2) proteins with abundances that decreased over the dilution series. (B) Beeswarm-boxplot of protein standard and complex protein sample fold changes. (C) Frequency distribution plot of CVs of complex protein sample.

For spectral count data, the majority of proteins had zero spectral counts at 1/16 and 1/32 dilutions; as such, protein abundance could only be assessed from dilutions 1 to 1/8. Spectral count data were further filtered such that at least two data points were present for every protein and analysed by hierarchical clustering (Fig. 8A). One distinct cluster was identified in which protein abundance did not change upon dilution, and protein standards were found in this cluster (see cluster marked with * in Fig. 8A). Although the median fold change of the proteins from the protein standard was 1.01-fold and agreed with expected fold change of 1-fold, the median fold change of the complex protein mix was 1.56-fold and underestimated the expected fold change of two (Fig. 8B). To determine the precision of spectral counting, CVs were calculated for quantified proteins. At zero dilution, 1/2 and 1/4 dilutions, the majority of proteins (~74%, 55% and 50%, respectively) were within a 20% CV threshold; in contrast, at 1/8 dilution, only a minority of proteins (~19%) were within a 20% CV threshold (Fig. 8C). These results indicate that while spectral counting could measure the abundance changes of a large number of proteins in a moderately complex IAC protein mix, the number of quantifiable proteins was reduced, fold changes underestimated, and the precision of readings was reduced compared to LC-MS ion intensity-based measurements of protein abundance.

**Fig 8.**
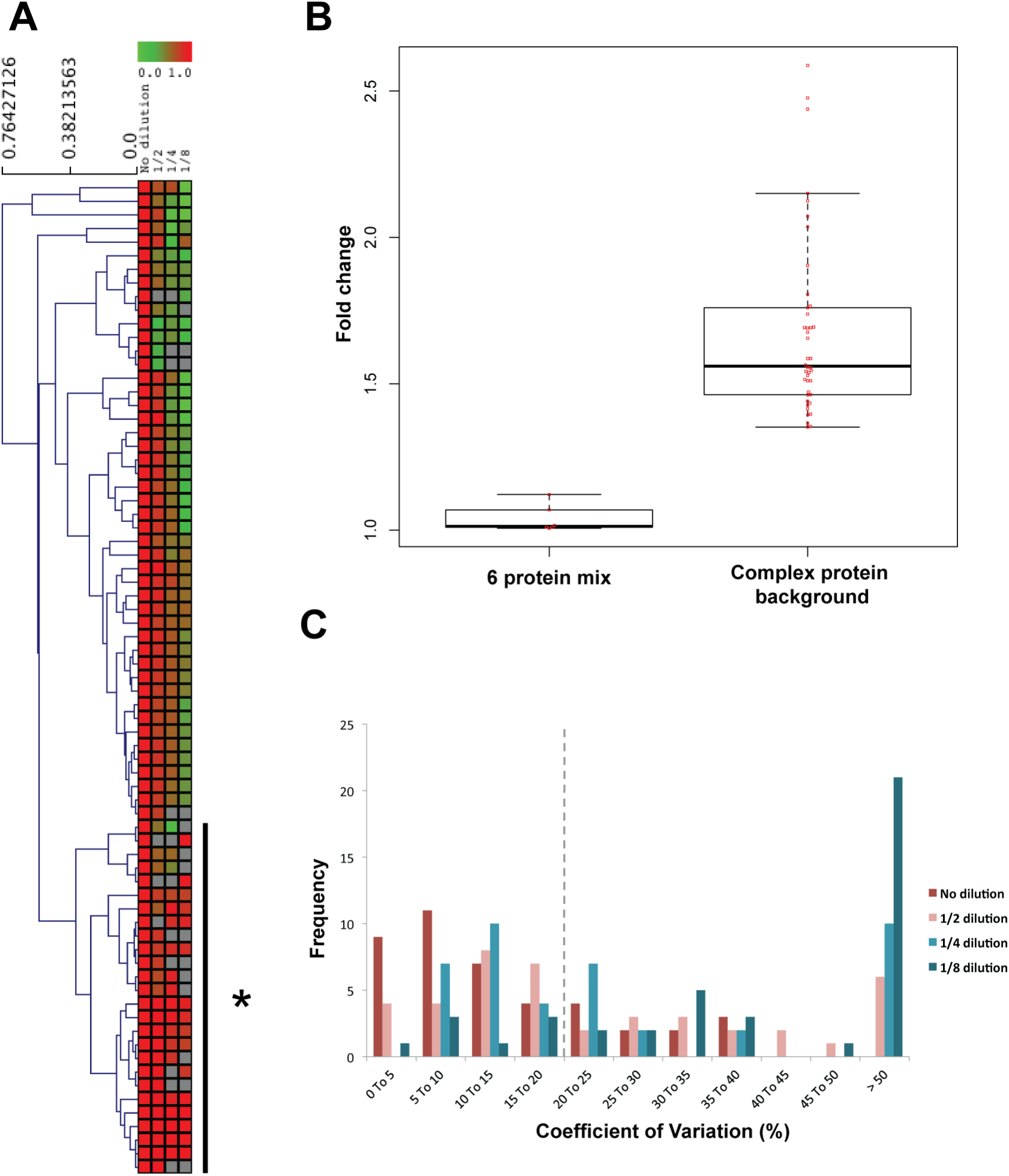
Quantitative analysis of changes to a complex protein mixture dilution series by spectral counting. (A) Heat map and dendrogram displaying the Euclidean distance-based hierarchical clustering of the scaled log_10_(average spectral counts) over four dilutions (no dilution to 1/8 dilution). The asterisk (*) denotes the cluster containing protein abundances that did not change over the dilution series. (B) Beeswarm-boxplot of protein standard and complex protein sample fold changes. (C) Frequency distribution plot of CVs of complex protein back-

## Discussion

In many LC-MS ion intensity quantification workflows, chromatographic alignment forms an integral part of the workflow [8,16]; however, although several groups have examined chromatographic alignment from an algorithmic perspective, few studies have addressed the impact of chromatographic alignment on protein quantification of biologically relevant samples. In addition, due to the improvements in mass spectrometers and analysis software, it is important to reassess and compare the label-free protein quantification strategies, spectral counting and LC-MS ion intensity quantification. Our major findings are that: 1) the quality of chromatographic alignment correlates with the precision of protein quantification, 2) a simple batch analysis strategy improves chromatographic alignment and 3) LC-MS ion intensity-based quantification is more accurate and precise than spectral counting in experiments where relatively few or a larger number of proteins were changing simultaneously.

Using technical replicates analysis of IAC samples with a range of Progenesis QI LC-MS alignment scores, we evaluated how chromatographic alignment quality affected protein quantification. We found that when chromatograms are not well aligned, precision of protein quantification was negatively affected, and the number of proteins identified in one condition and not another increased. Poor alignment of chromatograms negatively affected protein quantification due to conflicts in identifying misaligned features, inconsistencies in peak picking across conditions compared, and global normalisation factors being miscalculated. Therefore, confidence in the quantification process is intricately linked to the how well chromatograms are aligned.

Chromatographic drifts that arise due to differences in the performance of the chromatographic separation process can occur due to a host of different factors, including experimental variation, sample stability, temperature and pressure fluctuations, and changes in the chemical environment of the LC column due to aging, deposit build-up or interaction with different analytes in the sample [16,17,25,26]. To improve chromatographic alignment, warping algorithms have been developed to reduce both linear and non-linear chromatographic drifts [8,16,17,19]. However, many of these algorithms assume that the order of the peptides eluting from the chromatographic separation is the same [8,16,17]. Unfortunately, in many instances, the elution profile of a chromatographic run is not the same between runs and alignment algorithms are unable to deal with chromatographic alignment effectively. Indeed, we found that although well-aligned technical replicates displayed chromatographic drifts, the elution profile was mostly the same; instead, all technical replicates that displayed poor alignment had gross changes in the elution profile. It was observed that, in general, samples analysed close together in time in the same sample batch aligned better than samples analysed far apart in time and in different sample batches. Based on this observation, we developed a sample batch-running strategy to minimise chromatographic drifts and improve alignment (Fig 2). These findings are not only relevant to label-free quantification by Progenesis QI but will be useful to other LC-MS ion intensity quantification workflows that rely on chromatographic alignment.

Having established LC-MS ion intensity quantification dependence on chromatographic alignment, we tested its ability to quantify proteins accurately and precisely, and relate this to another label-free relative quantification strategy, spectral counting. Two experimental set-ups were used to mimic biologically relevant conditions: 1) few proteins changing in a moderately complex protein background, and 2) simultaneous changes to a large number of proteins (Fig. 3). Results from both experiments were complementary and showed that LC-MS ion intensity protein abundance measurements outperformed spectral counting in terms of accuracy and precision, and supported recent studies comparing both label-free methods in the estimation of absolute protein abundances [5,7, 27-29]. Moreover, although both label-free quantification methods responded linearly to increasing protein concentration, protein quantification measurements changed at almost the same rate for LC-MS ion intensity quantification but not spectral counting, suggesting that LC-MS ion intensity quantification is fairly robust to differences in physicochemical properties of peptides, number of peptides per proteins, and ionisation efficiencies of peptides. On the other hand, these factors, compounded by the intrinsic bias of DDA in spectral counting towards the most abundant proteins, probably result in the underestimation of the fold changes for proteins for spectral counting-based protein quantification. Therefore, we conclude for samples of moderate complexity, such as integrin adhesion complexes (IACs) [21-23], that spectral counting can provide useful estimates of relative protein abundance between samples; however, LC-MS ion intensity quantification was superior in terms of accuracy and precision.

To summarise, we have demonstrated that chromatographic alignment affects the quality of LC-MS ion intensity quantification, and developed a method to improve chromatographic alignment. Using this method, we designed experiments to test protein quantification by LC-MS ion intensity and compared it to spectral counting. In general, we found that when chromatographic alignment is good, protein quantification by LC-MS ion intensity outperforms spectral counting in terms of accuracy and precision.

## Acknowledgements

This work was supported by Cancer Research UK (grant C13329/A21671 to M.J.H.), the Wellcome Trust (grant 092015 to M.J.H.), a Wellcome Trust Institutional Strategic Support Fund award (grant 097820 to the University of Manchester) and a Wellcome Trust Four-Year PhD studentship (D.H.J.N). The mass spectrometers used in this study were purchased with grants from the Wellcome Trust and the University of Manchester Strategic Fund.

